# Enrichment of a CD4^-^CD8^-^ NK-like cytotoxic Vδ1/3 T cell subset in tuberculosis disease

**DOI:** 10.1101/2025.11.12.688134

**Authors:** Kendall Kearns, Rashmi Tippalagama, Ashu Chawla, Jason Greenbaum, Aruna Dharshan De Silva, Wathsala Gunasinghe, Judy Perera, Hansani Gunasekera, Darsha D. Senevirathne, Thomas J. Scriba, Julie G. Burel, Cecilia S. Lindestam Arlehamn, Bjoern Peters

## Abstract

Tuberculosis (TB), caused by *Mycobacterium tuberculosis*, remains a leading cause of global morbidity and mortality. Although gamma-delta (γδ) T cells have increasingly been suggested to contribute to the TB immune response, quantitative and qualitative differences in this immune cell compartment between healthy and TB diseased individuals are not well-characterized. In this study, we used single-cell RNA sequencing to provide a high-resolution characterization of CD4^-^CD8^-^ γδ T cells in peripheral blood across healthy *Mtb*-non-sensitized, healthy *Mtb*-sensitized, and TB disease pre-/post-treatment cohorts. We found upregulation of an activated and cytotoxic gene signature in γδ T cells of TB disease compared to both healthy cohorts. Strikingly, these differences persisted through one year following diagnosis of TB disease (corresponding to six months after completion of anti-TB therapy). We found that these transcriptomic differences were largely mediated by an NK-like cytotoxic Vδ1 and Vδ3 subset that was enriched in TB disease, with a unique Vδ3 TCR gene usage. Our findings suggest long-lasting changes in the CD4^-^CD8^-^ γδ T cell compartment and highlight Vδ3 cells, a previously underappreciated γδ T cell subset, as potentially important in TB.

## INTRODUCTION

Tuberculosis (TB), an airborne infection caused by *Mycobacterium tuberculosis* (*Mtb*), is reported to affect over 10 million people annually and remains a leading cause of death worldwide, despite the availability of antibiotic treatments and the Bacillus Calmette-Guérin (BCG) vaccine (1). Conventional alpha-beta (αβ) CD4 T cells recognizing peptide antigens are well-established as the primary lymphocyte subset involved in protective immunity to TB (2). However, growing evidence supports a role for unconventional T cells in the TB immune response. In our previous studies aimed at identifying peptide epitopes in *Mtb* (3,4), we noted IFNγ production in CD4^-^CD8^-^ T cells stimulated with *Mtb* extracts that was not present after *Mtb*-derived peptide stimulation. Here, we wanted to characterize the CD4^-^CD8^-^ T cell population, of which γδ T cells comprise a major proportion, in response to stimulation with *Mtb* antigens and *ex vivo* at the transcriptomic level across TB cohorts.

γδ T cells differ from conventional αβ T cells in that their T cell receptors (TCR) are encoded by a different set of genes and they are largely CD4^-^CD8^-^ instead of either CD4^+^ or CD8^+^ (5). They have been reported to be able to respond to non-peptide antigens and may not be MHC-restricted (6,7). They also hold features of both innate and adaptive immune cells, combining rapid responses with specificity, poising γδ T cells as potentially important in early stages of *Mtb* infection. The two predominant γδ T cell subsets are Vδ1 and Vδ2, which are named by their expression of *TRDV1* and *TRDV2* TCR genes, respectively, though other smaller subsets like Vδ3 (expressing *TRDV3*) exist (8). Vδ2 cells are the most prominent in circulation, of which >90% have Vγ9Vδ2 TCR pairing (9). Vγ9Vδ2 cells are known to recognize microbe-derived phosphoantigens, like HMBPP, that are present in *Mtb* and BCG (10). Vδ1 cells are less common in the periphery but are found at higher frequencies in tissues such as the mucosa (9). Vδ1 cells have been found to be enriched in the lungs of individuals with TB disease, suggesting they may be poised to quickly respond to *Mtb* challenge (11). Similarly, Vδ3 cells are rarely detected in circulation but can be found at higher frequencies in tissues like the liver and gut (8).

As Vδ2 cells are the most prominent γδ T cell subset in the circulation, they are the most well-studied in the context of human TB infection. Interestingly, several studies have demonstrated a significant decrease in the Vδ2:Vδ1 ratio in the circulation of individuals with TB disease compared to healthy individuals without evidence for TB infection (11,12) as well as in the lungs compared to individuals with sarcoidosis (13), suggesting potential induction of activation-induced cell death of Vδ2 T cells or expansion/migration of Vδ1 T cells during TB infection. In terms of functionality, γδ T cells can expand and produce IFNγ following BCG vaccination, provide memory-like responses, and protect against TB in nonhuman primate models (14–17). γδ T cells also exhibit cytotoxic and regulatory functions, which may be involved in protection against TB (15,18). As these functions are critical in responding to TB infection, deeper characterization of γδ T cells would be beneficial in better understanding their roles in the TB immune response across disease states and anti-TB treatment.

Currently, there is an overall lack of comprehensive data on how γδ T cells and their phenotypes differ between TB cohorts and how they are affected by TB treatment. Several studies have analyzed γδ T cells with single-cell RNA sequencing (scRNAseq) in the context of TB, highlighting trained immunity and subset heterogeneity of γδ T cells (15,19,20). However, these studies were limited in cell numbers (< 7,000 cells per study) and did not examine transcriptomic changes after starting anti-TB therapy. To build on this and further investigate the reactivity of CD4^-^CD8^-^ T cells across TB cohorts, we utilized blood samples from two healthy cohorts with or without *Mtb* sensitization and one TB disease cohort with longitudinal sampling at four timepoints pre- and post-treatment. We quantified IFNγ production by CD4^-^CD8^-^ T cells in response to different *Mtb* antigens and identified γδ T cells as the major source of IFNγ. We then isolated over 19,000 γδ T cells for scRNAseq analysis to further probe their transcriptomic profiles across TB cohorts. We hypothesized that this comprehensive dataset could help identify specific subsets of γδ T cells associated with TB disease compared to healthy cohorts with and without TB exposure as well as transcriptomic changes throughout anti-TB treatment.

## RESULTS

### CD4^-^CD8^-^ γδ T cells respond to BCG but not an *Mtb*-derived peptide pool

To investigate reactivity to *Mtb* antigens across T cell populations, we compared IFNγ production in CD4^+^, CD8^+^, and CD4^-^CD8^-^ T cell subsets in response to *M. bovis* BCG and *Mtb*-derived peptide epitope pool (MTB300) stimulation (3). We isolated peripheral blood mononuclear cells (PBMC) from individuals who were healthy *Mtb*-sensitized (determined by a positivity to IFNγ release assay, IGRA+) or those who had TB disease (active TB, ATB). PBMC samples were stimulated with either BCG or MTB300 and IFNγ production was measured after 12 hours (**Fig. S1**). In the IGRA+ cohort, CD4^+^ T cells produced significantly more IFNγ than CD8^+^ T cells in response to both BCG and MTB300 stimulation (**Fig. 1A**), supporting previous findings (21,22). We also found that CD4^-^CD8^-^ T cells produced more IFNγ than CD8^+^ T cells when stimulated with BCG but not MTB300 (**Fig. 1A**, left), which was also observed in the TB disease cohort (**Fig. 1A**, right). Characterization of the CD4^-^CD8^-^IFNγ^+^ population of both IGRA+ and ATB cohorts via flow cytometry revealed that it was predominantly γδ T cells and not MAIT cells (Vα7.2^+^CD161^+^) or other γδTCR^-^ subsets (**Fig. 1B**). This is in line with previous reports indicating that γδ T cells produce IFNγ in response to *Mtb* stimulation (14,16). These findings prompted further investigation of CD4^-^CD8^-^ γδ T cells in the context of TB.

**Figure 1.**
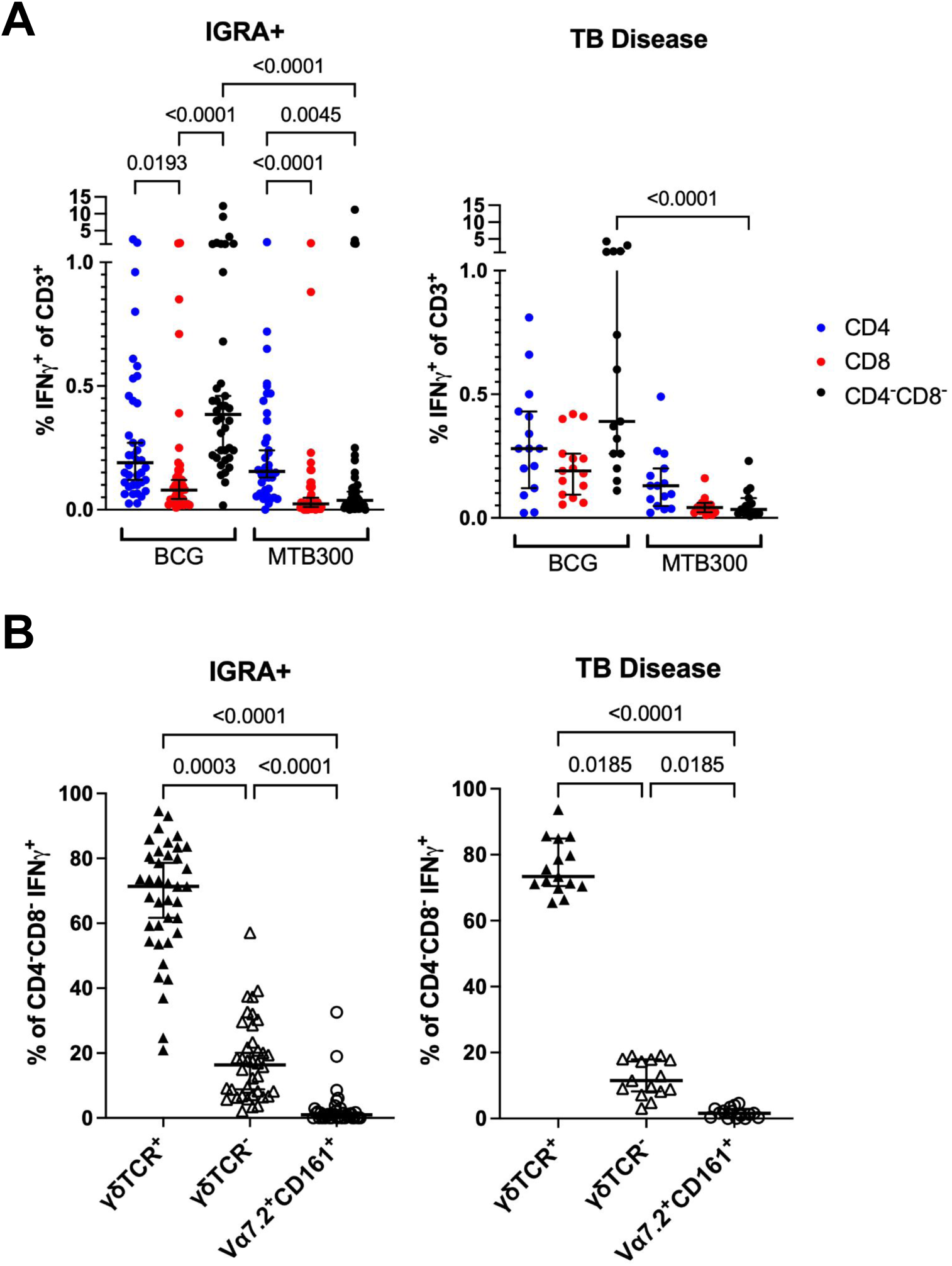
CD4^-^CD8^-^ γδ T cells produce IFNγ in response to BCG stimulation. A) Percent of IFNγ^+^ cells within the CD3^+^ population after stimulation with either BCG or MTB300 peptide pool separated by CD4^+^ (blue), CD8^+^ (red), and CD4^-^CD8^-^ (black) subsets. Samples isolated from IGRA+ and TB diseased individuals were utilized. Friedman test with Dunn’s multiple comparisons test. B) Percent of CD4^-^CD8^-^IFNγ^+^ cells stimulated with BCG that were γδTCR^+^, γδTCR^-^, or Vɑ7.2^+^CD161^+^ (MAIT cells) within the IGRA+ and TB disease cohorts. Friedman test with Dunn’s multiple comparisons test.

### CD4^-^CD8^-^ γδ T cell frequencies do not change across TB cohorts and timepoints

To further investigate CD4^-^CD8^-^ γδ T cells, we measured their frequency using flow cytometry in PBMCs isolated from IGRA+ individuals and individuals with ATB across four timepoints throughout anti-TB treatment in addition to healthy *Mtb*-non-sensitized controls (IGRA-) (**Fig. 2A**). We observed no significant differences in the frequencies of CD4^-^CD8^-^ γδ T cells across the cohorts and during anti-TB treatment (**Fig. S2A**) as well as between males and females (**Fig. S2B**). Additionally, there were no significant differences in CD4^+^ and CD8^+^ γδ T cell population frequencies (**Fig. S2C**). We also noted high variability across individuals within each cohort which may be indicative of participant-intrinsic factors that influence γδ T cell frequencies (4,5). Thus, there were no differences in the frequency of total CD4^-^CD8^-^ γδ T cells across cohorts.

**Figure 2.**
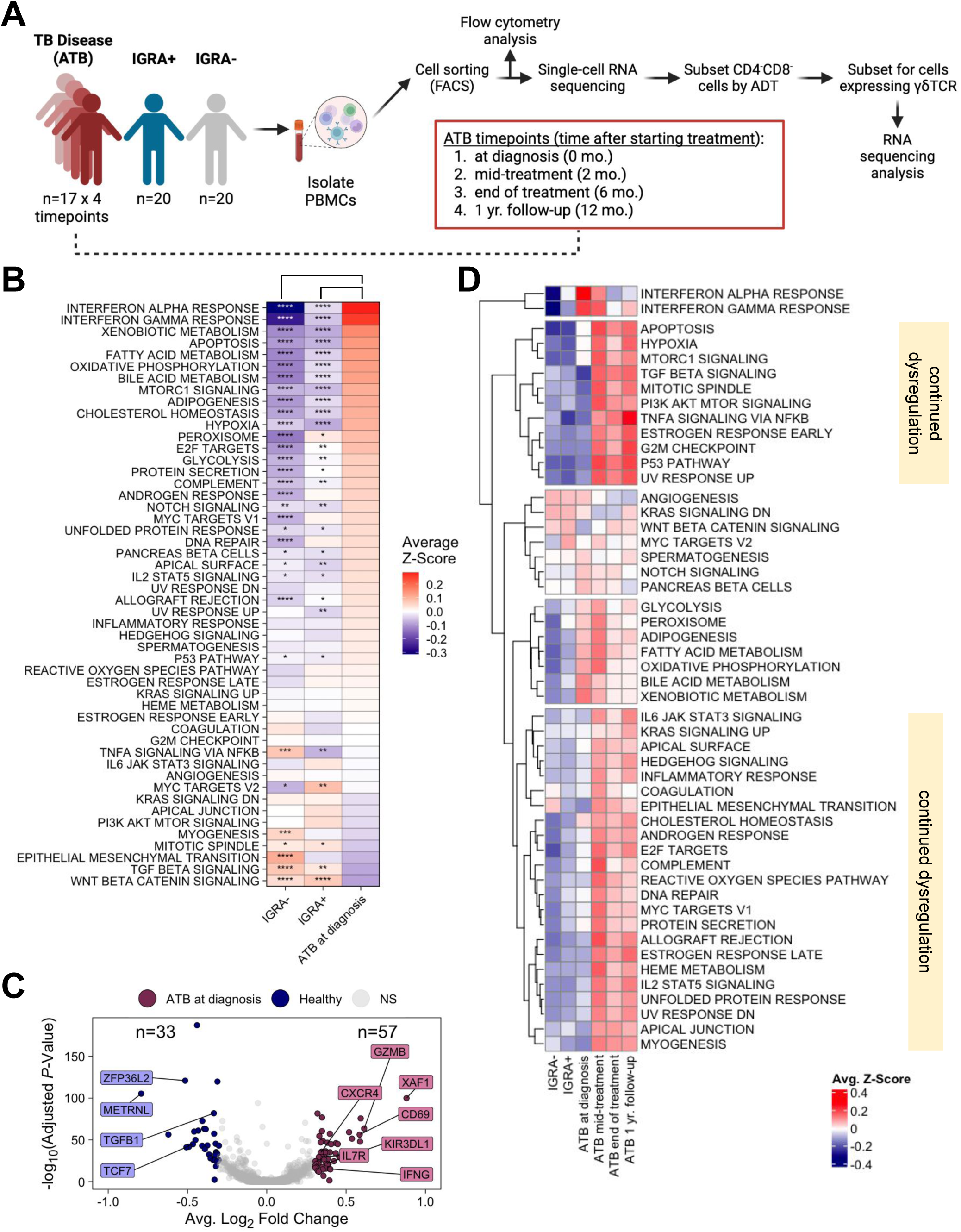
**CD4^-^CD8^-^ γδ T cells from ATB and healthy cohorts are transcriptionally distinct.** A) Experimental outline for examining CD4^-^CD8^-^ γδ T cells. Figure created with BioRender. B) Enrichment of hallmark gene sets within the IGRA-, IGRA+, and ATB at diagnosis cohorts. Enrichment scores were calculated for each cell, scaled for each gene set to obtain Z-scores, and the average Z-score (plotted) was calculated per pathway within each cohort. The pathways are ordered by the difference in ATB at diagnosis average Z-score and the average Z-score of IGRA- and IGRA+ cohorts. Statistical tests were performed using pairwise Wilcoxon rank-sum tests with Benjamini-Hochberg multiple testing correction of Z-scores between ATB at diagnosis and IGRA- or ATB at diagnosis and IGRA+ cohorts, with the significant adjusted *P*-values displayed in the IGRA- and IGRA+ columns, respectively. \**P* < 0.05; \*\**P* < 0.01; \*\*\**P* < 0.001; \*\*\*\**P* < 0.0001. C) Differential gene expression analysis between ATB at diagnosis (red) vs. IGRA-/IGRA+ (healthy, blue). Labeled genes include the gene with the highest fold change and genes selected based on biological relevance. D) Average GSEA Z-scores for all cohorts/timepoints.

### CD4^-^CD8^-^ γδ T cells from ATB and healthy cohorts are transcriptionally distinct

To identify differences in transcriptional phenotypes between CD4^-^CD8^-^ γδ T cells isolated from individuals across ATB and healthy cohorts, we performed single-cell RNA sequencing (scRNAseq) on *ex vivo* fluorescence-activated sorted CD8^-^ T cells (**Fig. 2A**, **Fig. S3**). Following scRNAseq, we filtered for cells that were CD4^-^ by antibody-derived tag (ADT), passed quality control metrics, and expressed γδTCR genes (Methods, **Fig. S4A-B**), which resulted in 19,807 CD4^-^CD8^-^ γδ T cells used for downstream analyses.

First, we performed single-cell gene set enrichment analysis (GSEA) with hallmark gene sets using the escape (v2.0.0) R package (25) to identify global differences in CD4^-^CD8^-^ γδ T cells between the IGRA-, IGRA+, and ATB at diagnosis cohorts. We first focused on the ATB at diagnosis cohort, as it is the most acute timepoint and was expected to have the most differences in comparison to the healthy cohorts. Indeed, the pathway analysis showed positively correlated changes in the two healthy cohorts, while both were negative correlated with the ATB cohort (**Fig. S5A**), suggesting that the IGRA- and IGRA+ cohorts are transcriptionally similar. Cells isolated from PBMC samples of the ATB at diagnosis cohort were significantly enriched for several immune activation pathways, including interferon alpha/gamma response and several metabolic pathways, compared to both IGRA- and IGRA+ cohorts (**Fig. 2B**).

Since GSEA showed that the hallmark pathways were positively correlated between IGRA- and IGRA+ cohorts (**Fig. S5A**), we performed differential gene expression analysis to directly compare the two cohorts. This revealed a total of only five differentially expressed genes between the IGRA- and IGRA+ cohorts (**Fig. S5B**), further suggesting that these two groups were very similar based on transcriptional profiles. Thus, we combined the IGRA- and IGRA+ cohorts into a single “healthy” cohort for subsequent analysis. Using this combined healthy cohort, we performed a differential gene expression analysis with the ATB cohort. There were 33 and 57 significantly upregulated genes (DEGs) in the combined healthy cohort and ATB at diagnosis cohort, respectively (**Fig. 2C**, **Supp.** **Table 1**). Healthy cohort upregulated DEGs included *TCF7*, *TGFB1*, *ZFP36L2*, and *METRNL*, which are associated with a naïve and inhibitive phenotype. In the ATB at diagnosis cohort, DEGs were associated with an activated/cytotoxic phenotype (*IFNG*, *GZMB*, *CD69*) and migration (*CXCR4*). These data suggest that TB disease significantly affects the transcriptomic profiles of circulating CD4^-^CD8^-^ γδ T cells, while those from healthy *Mtb*-sensitized and non-sensitized individuals are similar to each other.

#### Long-lasting transcriptomic changes are observed post-anti-TB treatment

Next, we sought to determine whether the transcriptomic profiles of ATB samples collected after anti-TB treatment (mid-treatment (∼2 month), end of treatment (∼6 month) and post-treatment/1 yr. follow-up (∼12 month)) would gradually return to those of healthy samples. Indeed, for the interferon alpha/gamma response pathways again enriched in ATB at diagnosis (**Fig. 2D**), we found a lower enrichment mid-treatment which continued to decrease through the 1-year follow-up of ATB, suggesting that treatment reduced the amount of interferon alpha/gamma that CD4^-^CD8^-^ γδ T cells respond to. However, other pathways showed a continued dysregulation in samples collected after beginning anti-TB therapy compared to healthy cohorts and ATB at diagnosis (**Fig. 2D**). Some of these pathways included apoptosis, reactive oxygen species pathway, and unfolded protein response, potentially suggesting a contraction phase following reduction of bacterial burden. 73 genes were upregulated in ATB 1 yr. follow-up compared to IGRA-, some of which were also upregulated in ATB at diagnosis (*IFNG*, *CXCR4*, *CD69*), suggesting the activated transcriptomic profile persists six months after completing treatment (**Fig. S5C**). Altogether, these data suggest that active *Mtb* infection and/or anti-TB therapy induces significant, long-lasting transcriptomic changes aligned with an activated/cytotoxic phenotype compared to healthy cohorts.

#### Distinct γδ T cell clusters identified with scRNAseq

To investigate heterogeneity within CD4^-^CD8^-^ γδ T cells, we performed unbiased clustering based on gene expression, which revealed 10 clusters (**Fig. 3A**). The top-expressed genes were largely unique to each cluster (**Fig. 3B**, **Supp.** >**Table 1**). To annotate each cluster, we chose genes based on the gene with the highest average log_2_ fold change and other genes that had biological relevance per cluster as informed by Enrichr (26–28) and manual identification (**Fig. 3C**). Cluster 9 (*MALAT1*^+^ Vδ1/2) was determined to contain cells with low quality transcriptomes based on higher expression of mitochondrial genes and lower numbers of features and RNA counts (**Fig. S6A**) and was removed from downstream analyses.

**Figure 3.**
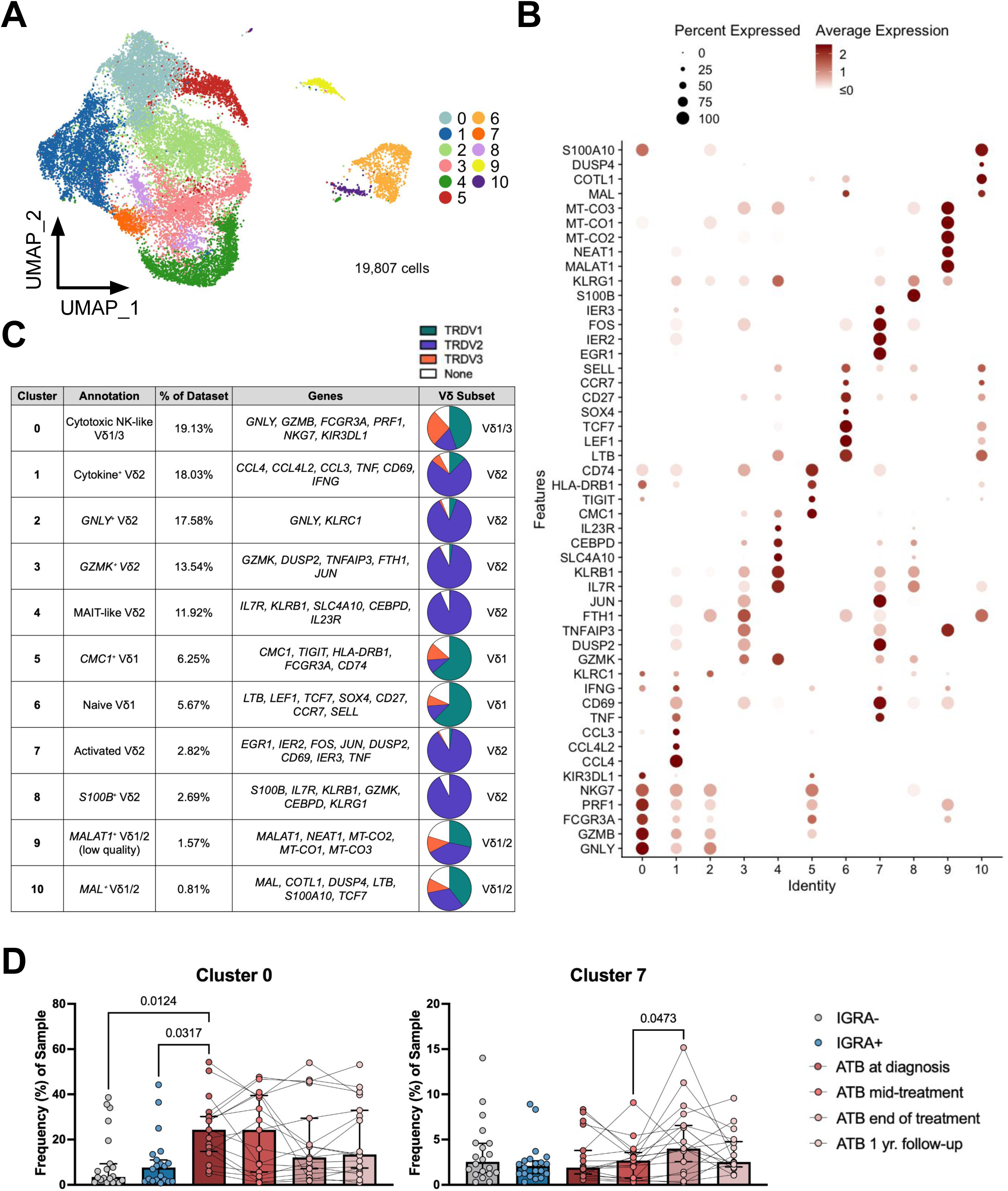
**scRNAseq revealed distinct clusters of CD4^-^CD8^-^ γδ T cells.** A) UMAP visualization of scRNA sequencing data colored by cluster. B) Top expressed genes of each cluster (y) with the average expression and percent of cells expressing each gene per cluster (x). C) Top genes, annotation, percent of dataset, Vδ subset, and proportion of cluster expressing *TRDV1*, *TRDV2*, or *TRDV3* (pie chart) for each cluster. A cell was determined to be *TRDV1*-, *TRDV2*-, or *TRDV3*-expressing if it had positive expression for one of the three genes and expression of 0 for the other two. A cluster was determined to be Vδ1, Vδ2, and/or Vδ3 if the associated gene was present in at least 25% of the cells within that cluster. D) Frequency of each sample found in clusters 0 and 7. For unpaired analyses, Kruskal-Wallis test with Dunn’s multiple comparisons test. For paired analyses, Friedman test with Dunn’s multiple comparisons test.

We assigned a Vδ subset category to each cluster based on the proportion of cells expressing *TRDV1*, *TRDV2*, or *TRDV3* in each cluster (**Fig. 3C**). The clusters were predominantly composed of either Vδ1 or Vδ2 cells, supporting previous findings that suggest the γδ T cell subsets are distinct populations (29,30). Additionally, most clusters were comprised of Vδ2 cells, which was expected as Vδ2 cells are the most prominent γδ T cell subset in circulation (9). Vδ2 clusters included cytokine^+^ Vδ2 (cluster 1), *GNLY*^+^ Vδ2 (cluster 2), *GZMK*^+^ Vδ2 (cluster 3), MAIT-like Vδ2 (cluster 4), activated Vδ2 (cluster 7), and *S100B*^+^ Vδ (cluster 8). Vδ1 clusters included *CMC1*^+^ Vδ1 (cluster 5) and naïve Vδ1 (cluster 5). Several mixed clusters were also identified: cytotoxic NK-like Vδ1/3 (cluster 0), *MALAT1*^+^ Vδ1/2 (cluster 9), and *MAL*^+^ Vδ1/2 (cluster 10).

Next, we calculated the cell cluster composition in each sample and compared the results between cohorts (**Fig. 3D**, **Fig. S6B**). Broadly, there were few significant differences, including across ATB timepoints. Cluster 7 showed a significant increase from ATB mid-treatment to end of treatment (**Fig. 3D**), indicating treatment-related differences. Cluster 0 was the only cluster that showed significant enrichment for ATB at diagnosis compared to both IGRA- and IGRA+ healthy cohorts (**Fig. 3D**). There were no significant longitudinal changes in cluster 0, though there was a trend for decreased frequencies through the 1 yr. follow-up timepoint. Overall, these data suggest potential involvement of cluster 0 (cytotoxic NK-like Vδ1/3) and cluster 7 (activated Vδ2) cell subsets in TB disease and resolution.

#### Several clusters exhibit high ATB module scores and *IFNG* expression

We next used the healthy vs. ATB at diagnosis DEGs (as reported in **Fig. 2C** and **Supp.** **Table 1**) to determine whether any clusters had an ATB-associated signature. We calculated ATB and healthy module scores for each cell using the Seurat (v4.4.0) R package (31), then obtained the log_2_-transformed ratio to represent the relative enrichment of ATB- and healthy-associated DEGs. We calculated adjusted module scores for each cluster (**Fig. 4A**). The clusters that exhibited enrichment (median > 0) of ATB-associated DEGs included clusters 0, 1, 2, 5, 7, and 8, which thus represented more activated populations compared to the rest of the clusters (**Fig. 3C**).

**Figure 4.**
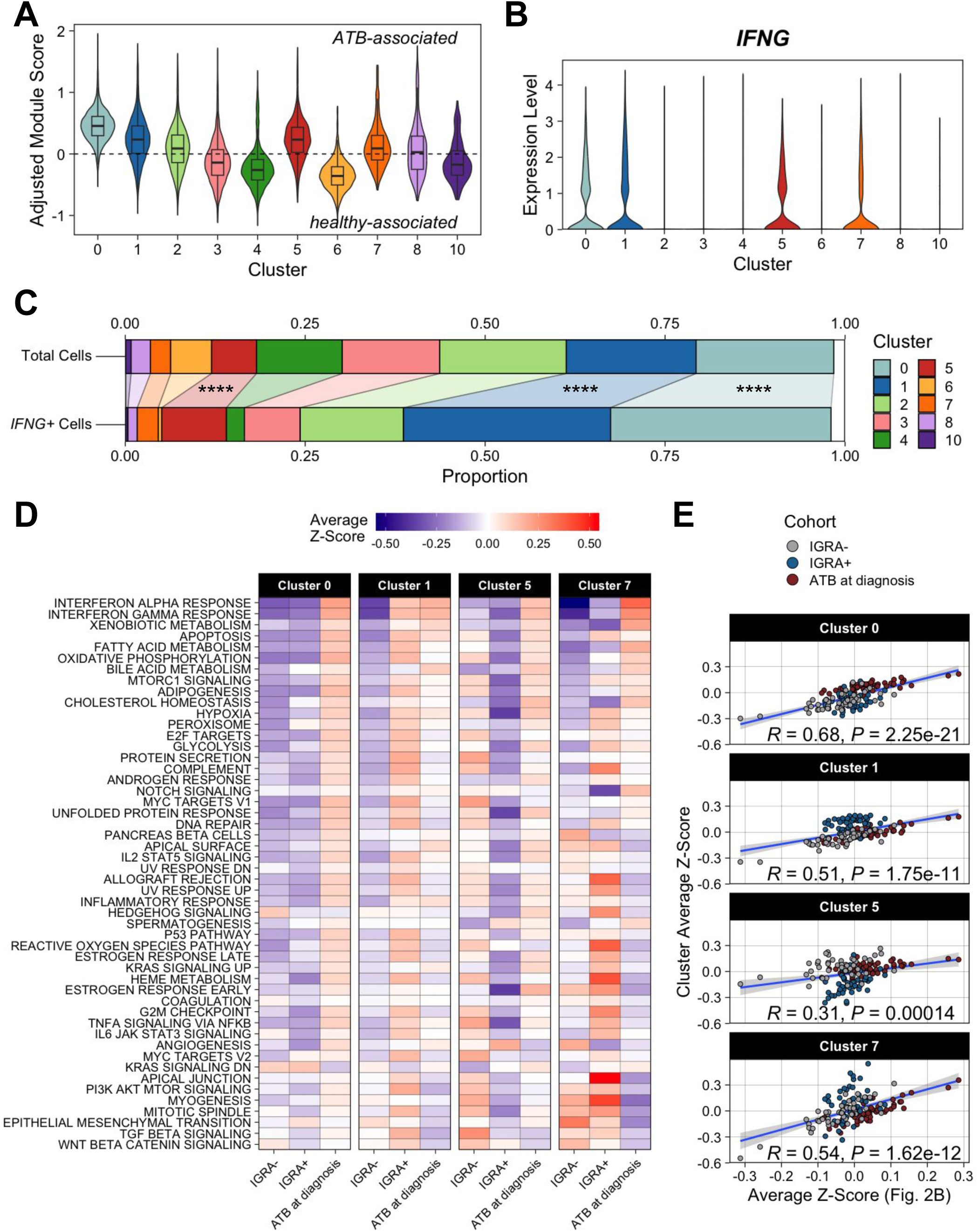
Several clusters exhibit high ATB module scores and *IFNG* expression. A) Module scores were calculated using the ATB- and healthy-associated genes in Fig. 2C. The adjusted score was calculated by dividing the ATB score + 1 by the healthy score + 1, then taking the log_2_ of the result. Cluster 9 was removed due to low quality. B) *IFNG* expression per cluster. Cluster 9 was removed due to low quality. C) Proportion of the total cell population (left) or of *IFNG*+ cells (right) that originate from each cluster. Cells with an *IFNG* expression level of greater than 1 were classified as *IFNG*+. Exact binomial test of whether the proportion of each cluster within the *IFNG*+ cell subset is significantly greater than the proportion of each cluster within the total cell population. \**P* < 0.05; \*\**P* < 0.01; \*\*\**P* < 0.001; \*\*\*\**P* < 0.0001. D) Enrichment of hallmark pathways within the IGRA-, IGRA+, and ATB at diagnosis cohorts for clusters 0, 1, 5, and 7. Pathways along the y axis are in the same order as in Fig. 2B. Enrichment scores were calculated for each cell and were scaled for each gene set to obtain Z-scores, which were then averaged within each cohort for each cluster. E) Plots showing the Pearson correlation between the average Z-scores from Fig. 2B (overall dataset) and from each cluster from the hallmark pathway enrichment analysis.

We then examined the expression of *IFNG* across clusters as it was found to be one of the ATB-associated DEGs (**Fig. 2C**), is known to be a critical cytokine involved in the TB immune response (32), and was produced by CD4^-^CD8^-^ γδ T cells in response to BCG stimulation (**Fig. 1**). Clusters 0, 1, 5, and 7 had high expression of *IFNG* (**Fig. 4B**), all of which also exhibited enrichment of ATB-associated DEGs (**Fig. 4A**). We calculated the proportion of *IFNG*^+^ cells (*IFNG* > 1) originating from each cluster and compared that to the proportion of each cluster within the total cell population (**Fig. 4C**). Clusters 0, 1, and 5 were found at significantly higher proportions within the *IFNG*^+^ population than expected within the total dataset, with over 50% of the *IFNG*^+^ cells originating from clusters 0 and 1.

Together, these data identify clusters 0, 1, 5, and 7 as the likely source of *IFNG*-producing CD4^-^CD8^-^ γδ T cells in ATB samples at diagnosis.

#### Cluster 0 is associated with enrichment in ATB at diagnosis

Next, we investigated whether differences across the cohorts existed in the four clusters with increased ATB module scores and *IFNG* expression to highlight subsets potentially involved in responding to TB disease. We performed single-cell GSEA for each of the four clusters with increased ATB module scores and *IFNG* expression. Clusters 1 and 5 each showed similar levels of enrichment between ATB at diagnosis and healthy cohorts, while the IGRA+ cohort in cluster 7 showed significant enrichment for many pathways compared to ATB at diagnosis (**Fig. 4D**). Although clusters 0, 5, and 7 exhibited high enrichment in interferon response pathways in ATB compared to healthy cohorts, suggesting these cells may be responding to infection, only cluster 0 exhibited an enrichment pattern similar to that of the global analysis of all CD4^-^CD8^-^ γδ T cells (**Fig. 2B**) with high enrichment of many of the same pathways in ATB at diagnosis compared to healthy cohorts (**Fig. 4D**). When we compared the average Z-scores of each pathway between each of the four clusters and the overall dataset, we found that cluster 0 had the strongest and most significant correlation (**Fig. 4E**), indicating an enrichment profile that most closely mirrors that of the overall dataset compared to the other clusters. Overall, these data suggest that cluster 0 is associated with enrichment in ATB at diagnosis in terms of pathway enrichment and may be involved in the anti-TB immune response.

### Vδ3 cells were significantly enriched in ATB at diagnosis

We next analyzed expression of TCR δ genes to determine if there was a preference for specific gene usage between the ATB and healthy cohorts. First, we calculated the Vδ2:Vδ1 ratio per sample using the expression level of *TRDV2* and *TRDV1* genes and found a trend for lower ratios in ATB compared to healthy cohorts that persisted through the last timepoint post-treatment (**Fig. 5A**), supporting previous findings (11,12). Consistent with this finding, we also noted an increase in the proportion of cells having nonzero expression of *TRDV1* and *TRDV3* and a corresponding decrease in cells expressing *TRDV2* across ATB cohorts compared to healthy individuals (**Fig. 5B**). To further validate these findings, we used flow cytometry to measure the frequency of γδ T cell subsets. Since there is no commercially available Vδ3 antibody, we used Vδ1^-^Vδ2^-^ cells within the γδ T cell population as a proxy for Vδ3 cells as they are the next most abundant γδ T cell subset after Vδ1 and Vδ2 cells (33). We found a significant increase in Vδ1^-^Vδ2^-^ cell frequency in ATB at diagnosis compared to both IGRA- and IGRA+ cohorts (**Fig. 5C**). We also found a trend for increased Vδ1 and significantly decreased Vδ2 cell frequencies in ATB at diagnosis compared to the IGRA- cohort (**Fig. 5C**).

**Figure 5.**
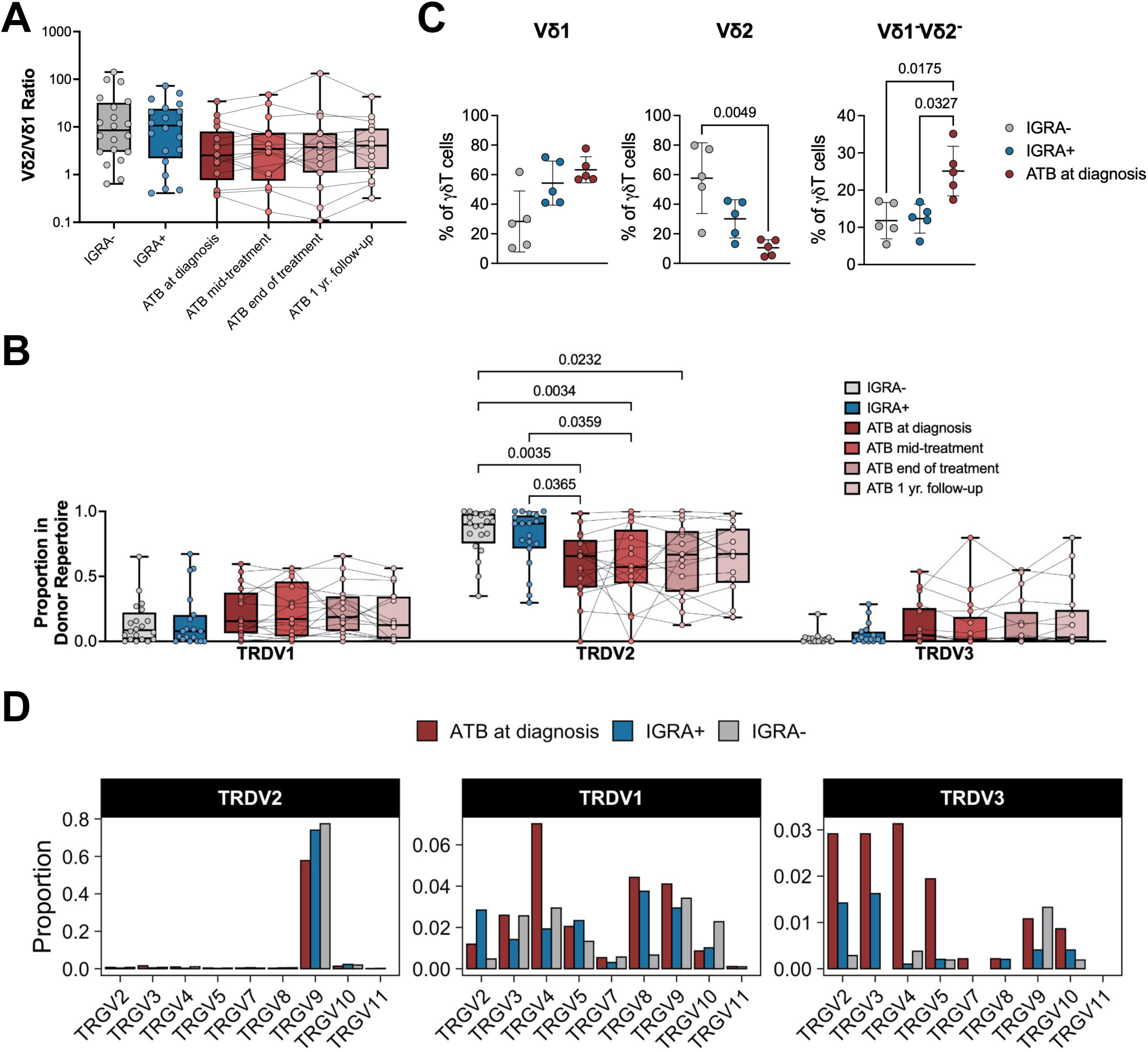
**Vδ3 TCR is significantly enriched in ATB at diagnosis.** A) Vδ2:Vδ1 ratio calculated for each cohort using expression of *TRDV1* and *TRDV2*. B) Proportion of the total donor TCR repertoire expressing *TRDV1*, *TRDV2*, or *TRDV3* in IGRA-, IGRA+, and ATB cohorts across treatment timepoints for each donor. Ordinary two-way ANOVA with Tukey’s multiple comparisons test. C) Frequency of Vδ1, Vδ2, and Vδ1^-^Vδ2^-^ T cells within the γδ T cell population in IGRA-, IGRA+, and ATB at diagnosis cohorts using flow cytometry. Kruskal-Wallis test with Dunn’s multiple comparisons test within each subset D) Proportion of each cohort with specific *TRGV* gene pairings for *TRDV1*, *TRDV2*, and *TRDV3*.

Next, we examined the usage of specific TCR gamma and delta genes pairs to potentially identify combinations associated with TB disease. We focused on the healthy and ATB at diagnosis cohorts, as these displayed the most striking differences in overall phenotypes. We utilized expression values of TCR variable (V) genes and only included cells expressing both TRGV and TRDV genes in this analysis. We found that the *TRDV2* TCRs nearly exclusively pair with *TRGV9* in all disease conditions, as previously described (Fig. 5D). For *TRDV1* and *TRDV3* TCRs, gamma chain usage was diverse and varied across the cohorts. The increase of *TRDV1* usage in ATB at diagnosis was associated specifically with the *TRGV4* gamma chain, which was also substantially increased in combination with *TRDV3* (Fig. 5D). This may indicate that *TRGV4*/*TRDV1* and *TRGV4*/*TRDV3* gene pairs may be involved in the anti-TB immune response (34). Overall, these data suggest that specific γδ TCR chain pairings are associated with TB disease.

## DISCUSSION

In this study, we characterized the *ex vivo* frequencies and transcriptional profiles of CD4^-^CD8^-^ γδ T cells across healthy and ATB cohorts and timepoints throughout anti-TB treatment. We found that γδ T cells were the primary producers of IFNγ in response to BCG (but not MTB300 peptide pool, as expected) within the CD4^-^CD8^-^ T cell population, supporting previous studies (14,16). Despite stable *ex vivo* frequencies across cohorts and timepoints, CD4^-^CD8^-^ γδ T cells exhibited strong transcriptional differences in ATB *ex vivo*, suggesting functional reprogramming and a bias towards specific γδ T cell subsets in response to *Mtb* infection and/or anti-TB treatment.

Using gene set enrichment analysis, we found that CD4^-^CD8^-^ γδ T cells from ATB patients were enriched for many immune activation and metabolic pathways. The enrichment of interferon alpha/gamma response pathways in ATB is consistent with previous findings confirming the involvement of these pathways in the TB immune response (35–38). The metabolic pathways (e.g., oxidative phosphorylation, glycolysis) that were also upregulated combined features of previously described IL-17-and IFNγ-producing γδ T cells, which are associated with tumor growth/metastasis and tumor surveillance, respectively (39–41). While the previous studies were focused on γδ T cells in cancer, IL-17 and IFNγ have also been implicated in TB immunity, of which γδ T cells may be an important source (2,42–45). Additionally, CD4^-^CD8^-^ γδ T cells from ATB showed increased expression of genes related to activation, cytotoxicity, and migration (*CD69*, *IFNG*, *GZMB*, *CXCR4*), suggesting that CD4^-^CD8^-^ γδ T cells with an effector phenotype and potential to migrate to the lung are involved in *Mtb* immune responses.

We identified four clusters (0, 1, 5, and 7) as clusters of interest due to their high ATB gene module scores and *IFNG* expression, suggesting that these subsets may be involved in anti-TB immunity. Cluster 0 (cytotoxic NK-like Vδ1/3) had the highest proportion of Vδ3 cells and was the only cluster that was significantly enriched in ATB at diagnosis compared to healthy cohorts. We also found a decreased Vδ2:Vδ1 ratio and increased frequency of Vδ3 cells in the circulation of TB patients. This was a striking observation, as Vδ1 and Vδ3 cells are more commonly found in tissues compared to circulation and may suggest a role for these subsets in TB (8,11,12). Other studies have revealed that circulating Vδ1 and Vδ3 cells increase in frequency in other contexts such as cancer and viral infections and have suggested that these cells may be involved in memory formation and antiviral immunity (46–48). Our findings indicate that these two Vδ subsets may also play a role in anti-TB immunity.

Additionally, cluster 0, which we annotated as a cytotoxic NK-like Vδ1/3 subset, closely resembles a previously defined CD8^+^ NK-like γδ T cell population in individuals with persistent *Mtb* infection (49). This population was found to consist mostly of *TRDV1*-expressing cells and exhibited significantly decreased ability for TCR-mediated activation with a strong capacity for CD16-mediated cytotoxicity. Thus, cluster 0 cells may also be activated through non-TCR-mediated mechanisms such as antibody-dependent cellular cytotoxicity.

We observed a decrease in the Vδ2:Vδ1 ratio in ATB compared to healthy cohorts, supporting previous findings (11,12). Along with this, there was an increase in *TRDV3* and decrease in *TRDV2* usage at the gene level across ATB timepoints and at the protein level in ATB at diagnosis in comparison to healthy cohorts. Further analysis revealed a diverse TCR gene pair repertoire. Only gene pairs including *TRDV3* or TRDV1 were found to be significantly associated with ATB via permutation test. Both *TRDV1* and *TRDV3* gene pairs associated with ATB included *TRGV4*, which has been found at higher frequencies in the blood of TB patients (34), suggesting potential involvement in the anti-TB immune response. Although Vδ3 cells are rarely found in the blood of healthy individuals, higher frequencies in the circulation have been reported in the context of cancer and viral infections (50). However, their profile in TB is not well-understood, and this work presents novel findings of Vδ3 T cells significantly associated with TB.

All four timepoints analyzed in the ATB cohort exhibited transcriptomic profiles distinct from that of healthy individuals. Additionally, the bias for Vδ1 and Vδ3 cells was present throughout all four timepoints. This is remarkable, as the fourth timepoint corresponds to six months after the end of anti-TB therapy, approximately one-year post-diagnosis. Together, these results suggest that *Mtb* infection and/or anti-TB treatment induce long-lasting transcriptomic changes in CD4^-^CD8^-^γδ T cells. Durable transcriptomic differences between healthy and ATB follow-up cohorts have been shown previously, though the number of DEGs gradually decreased compared to that of healthy vs. ATB at diagnosis and these data were obtained from whole blood (51). It remains to be defined whether these changes are only long-lasting in specific subsets, such as CD4^-^CD8^-^γδ T cells as described here, or systemic.

Our study has several limitations. Our analysis was restricted to circulating γδ T cells and does not directly assess the responses occurring in the lung during TB infection, which may be different. Additionally, whether and how the different γδ T cell clusters herein described recognize *Mtb* and how this may differ across TB disease states were not addressed by the present study. Functionally validating the effector potential of these subsets, particularly Vδ3 cells, to determine whether they contribute to host protection, immunopathology, or both would be crucial to better understanding the roles of γδ T cells in the *Mtb* immune response. Further work using targeted TCRγδ sequencing could reveal additional information on specific CDR3 sequences associated with TB disease and lead to the identification of putative ligands.

In conclusion, our study reveals long-lasting transcriptomic changes in CD4^-^CD8^-^ γδ T cells in ATB compared to healthy individuals. We also identify a distinct population of CD4^-^CD8^-^ γδ T cells, cytotoxic NK-like Vδ1/3 cells, that is enriched in ATB at diagnosis in terms of frequency, gene module scores, and pathway enrichment. Specifically, Vδ3 cells were significantly more associated with TB disease. These findings further contribute to our understanding of unconventional T cell responses in TB and highlight Vδ3 cells as a potentially important, yet understudied, component of the *Mtb* immune response.

## ACKNOWLEDGEMENTS

We thank the Flow Cytometry Core and Sequencing Core Facility at the La Jolla Institute for Immunology for their valued role in sample acquisition and sequencing. We acknowledge support from the National Institute of Health grant U19 AI118626 (to B.P.) and National Institutes of Health contract 75N93024C00057 (to B.P. and C.S.L.A).

## AUTHOR CONTRIBUTIONS

J.G.B., C.S.L.A, and B.P. participated in the design and direction of the study. K.K. and R.T. performed and analyzed the experiments. K.K. performed bioinformatic analysis. A.C., J.G., and R.T. assisted in bioinformatic analysis. R.T., J.P., H.G., D.D.S., W.G., A.D.DS., and T.J.S. recruited participants, performed clinical evaluations, and isolated PBMCs. K.K., J.B., C.S.L.A., and B.P. wrote the manuscript. All authors read, edited, and approved the manuscript before submission.

## MATERIALS & METHODS

### Ethics Statement

Human study participants were enrolled at the National Hospital for Respiratory Diseases, Welisara (Sri Lanka) or the South African Tuberculosis Vaccine Initiative, Western Cape Province (South Africa). Ethical approval was maintained through the La Jolla Institute for Immunology Institutional Review Board (IRB) or the Human Research Ethics Committee of the University of Cape Town. The University of Colombo Ethics Review Committee served as the National Institute of Health registered IRB for the Kotelawala Defence University. All clinical investigations were conducted according to the principles expressed in the Declaration of Helsinki, and all participants provided written informed consent prior to participation in the study.

### Participants and Samples

Participants were from South Africa and Sri Lanka and were age- and sex-matched across the sites. The individuals were segregated into three main cohorts: active TB (ATB), IGRA+, and IGRA-. The healthy IGRA+ cohort was identified as positive for the IFNγ-release assay (IGRA; QuantiFERON-TB Gold In-Tube, Cellestis or T-SPOT.TB, Oxford Immunotec) with the absence of symptoms consistent with TB or other clinical or radiographic signs of ATB. ATB was defined, in the absence of any other significant co-morbidity, as 1) presence of clinical symptoms and/or radiological/histological evidence of pulmonary TB and 2) microbiologically confirmed by sputum or *Mtb* culture. IGRA- individuals had no past medical history of TB or exposure to *Mtb* or evidence of *Mtb*-sensitization as confirmed by a negative IFNγ-release assay. All participants were confirmed negative for HIV. PBMCs were obtained by density gradient centrifugation (Ficoll-Hypaque, Amersham Biosciences/GE Healthcare) from leukapheresis or whole-blood samples, according to the manufacturer’s instructions. Cells were resuspended at 50-100 million cells per milliliter in FBS (Gemini Bio-Products) containing 10% DMSO (Sigma) and cryopreserved in liquid nitrogen.

## PBMC Thawing

Cryopreserved PBMCs were quickly thawed by incubating each cryovial at 37°C for 1-2 minutes. Cells were then transferred into 9 mL of cold HR5 medium (RPMI 1640 with L-Glutamin and 25 mM HEPES [Omega Scientific], supplemented with 5% human AB serum [GemCell], 1% Penicillin/Streptomycin [Gibco], and 1% Glutamax [Gibco]) and 20 U/mL Benzonase Nuclease (Millipore). Cells were centrifuged and resuspended in medium to determine cell concentration and viability using trypan blue and a hemacytometer.

### Intracellular Cytokine Staining

1x10^6^ thawed PBMCs were plated per well in 96-well round-bottom plates in HR5 medium. Cells were stimulated with *M. bovis* BCG-Danish (BCG vaccine SSI lyophilized powder for injection resuspended in provided diluent and stored at 4°C, *M. bovis* BCG Danish strain 1331; gifted from R. Mortensen, SSI, Denmark) or *M. bovis* Pasteur (frozen stocks at -80°C in 7H9 media, thawed, washed and resuspended in PBS; gifted from D. Barber, NIH, USA) at 100µg/mL or 50 µg/mL (2-8 x 10^6^ bacteria/mL), or MTB300 (2 µg/mL per peptide); anti-CD28 (1 µg/mL) and anti-CD49d (1 µg/mL) were present in all stimuli preparations. The plate was incubated for 5 hours at 37°C in a 5% CO_2_ incubator. BFA and monensin were added to each well per manufacturer’s recommended protocol and the plate was further incubated at 37°C for an additional 7 hours and then kept at 4°C until staining. Cells were stained with a surface panel containing anti-CD3 Alexa Fluor 700 (BD Biosciences, 557943), anti-CD4 APC-eFluor780 (eBioscience, 47004942), anti-CD8 brilliant violet 650 (BV650; BioLegend, 301042), anti-CD161 APC (eBioscience, 17161942), anti-TCR Vα7.2 phycoerythrin (PE)-Cy7 (BioLegend, 351712), anti-pan γδTCR PE (BD Biosciences, 347907), anti-CCR7 PerCP-Cy5.5 (BioLegend, 353220), anti-CD45RA eFluor450 (eBioscience, 48045842), anti-CD14 V500 (BD Biosciences, 561391), anti-CD19 V500 (BD Biosciences, 561121), and fixable viability dye eFluor 506 (eBioscience, 65-0866-14). Cells were fixed with a 4% paraformaldehyde solution for 10 minutes, permeabilized for 10 min in saponin buffer (0.2 g saponin, 400 µL sodium acetate, 4 mL BSA, in PBS to 40 mL total volume) and blocked with 10% FBS for 5 min on ice. Samples were then stained with anti-IFNγ FITC (eBioscience, 11731982) in saponin buffer for 30min at room temperature and then washed. Samples were analyzed on a BD LSR II. The gating strategy is shown in Figure S1.

### Flow Cytometry for 10x Single-Cell RNA Sequencing

Freshly thawed PBMCs were incubated for 10 minutes with Zombie UV Fixable Viability Kit (BioLegend) following the manufacturer’s protocol. To neutralize and block unbound viability dye, 10% fetal bovine serum (FBS) in 1X PBS was added to the cells followed by a centrifugation step. FcR blocking reagent (BioLegend) at a 1:50 dilution from the stock reagent solution was added to the cells. Samples were further incubated for 10 minutes at room temperature. For oligonucleotide-conjugated antibody staining, 2 µL of each TotalSeq^TM^-C antibody was added per sample in a final staining volume not exceeding 100 µL in 1X PBS and cells were incubated at room temperature for 15 minutes. TotalSeq^TM^-C antibodies that were used included anti-HLA-DR (clone L243, barcode 0159) and anti-CD4 (clone SK3, barcode 0045), as well as several anti-CD45 or anti-CD298 plus anti-β2-microglobulin antibodies to identify each sample in downstream analyses.

Samples were spun down and excess unbound antibody was removed before proceeding to the next step. A cocktail of fluorochrome-conjugated antibodies, 10 µL BD Horizon^TM^ Brilliant Stain Buffer Plus (BD Biosciences), and 1X PBS were added to a final volume of 100 µL. The antibodies included in the cocktail were anti-CCR7 BB515 (BD Biosciences, clone 2-L1-A), anti-CD11c BV785 (BioLegend, clone 3.9), anti-CD123 PerCP-Cy5.5 (BioLegend, clone 6H6), anti-CD14 BV480 (BD Biosciences, clone M5E2), anti-CD16 BUV737 (BD Biosciences, clone 3G8), anti-CD19 BV605 (BioLegend, clone HIB19), anti-CD1c BV650 (BioLegend, clone L161), anti-CD20 BUV563 (BD Biosciences, clone 2H7), anti-CD3 BUV805 (BD Biosciences, clone UCHT1), anti-CD38 PE-Dazzle594 (BioLegend, clone HIT2), anti-CD4 APC-eF780 (eBiosciences, clone RPA-T4), anti-CD45 BUV395 (BD Biosciences, clone HI30), anti-CD45RA BV570 (BioLegend, clone HI100), anti-CD56 APC (BioLegend, clone 5.1H11), anti-CD71 PE-Cy5 (BD Biosciences, clone M-A712), anti-CD8 BUV495 (BD Biosciences, clone RPA-T8), anti-HLA-DR PE-Cy7 (BioLegend, clone LN3), anti-IgD BV750 (BD Biosciences, clone IA6-2), and anti-TCRγδ BV421 (BD Biosciences, clone 11F2). Cells were incubated for 15 minutes in the dark at room temperature. Cells were washed once in 10 mL of 1X PBS to remove all unbound antibody and stored at 4°C protected from light for up to 4 hours until flow cytometry acquisition. Live singlets CD3^+^CD19^-^CD14^-^CD8^-^ cells were sorted using a BD Symphony S6 sorter and sequenced using droplet-based 10x Genomics.

### Single-Cell RNA Sequencing Data Preparation

The number of cells sorted per sample ranged from 241 to 3,333 depending on the quality of each sample. Sorted cells were resuspended at a concentration of 1,800 cells per µL in ice-cold PBS with 0.04% BSA. Single-cell suspensions were then immediately loaded on the 10x Genomics Chromium Controller with a loading target of 30,000-60,000 cells. Libraries were generated using the Chromium Next GEM Single Cell 5’ Reagent Kit v2 (Dual Index) per the manufacturer’s instructions. Additional steps were included for the amplification of HTO barcodes and V(D)J libraries (10x Genomics). Libraries were sequenced on an Illumina NovaSeq with a target of 30,000 reads per cell RNA library, 5,000 reads per cell HTO barcode library, and 5,000 reads per cell for V(D)J libraries. As the V(D)J libraries utilized the default TCRαβ primers, the data was not used in this work but were analyzed further in another study.

### scRNA-Seq analysis – hashtag demultiplexing, quality control, clustering

Libraries were demultiplexed and FASTQ files were created using CellRanger *mkfastq* and mapped with the CellRanger *multi* pipeline (v5.0.0) (52) against reference genome GRCh38 (2020-A). TCR genes belonging to TCR α, β, γ, or δ from the GEX matrix were aggregated to prevent clusters forming based on individual TCR gene expression. The GEX and HTO/ADT data were aggregated to form a Seurat object using the R package Seurat (v4.4.0) (31) and quality control metrics (mitochondrial percent < 8, nFeature_RNA ≥ 1,000, nCount_RNA < 15,000, HTO singlets) and assay normalizations (*SCTransform* for GEX, CLT for HTO/ADT) were applied.

Cells within the Seurat object that had expression of both aggregated TCR γ and δ genes (SCT assay) (Fig. S4) and no expression of ADT-CD4 were selected to create a new Seurat object containing CD4^-^CD8^-^ γδ T cells. The new Seurat object was then subjected to dimensionality reduction and clustering using *FindNeighbors*, *RunUMAP* with 30 dimensions, and *FindClusters* with a resolution of 0.5. Cluster-specific markers were obtained using *PrepSCTFindMarkers* and *FindAllMarkers* with a min.pct of 0.25 and a logfc.threshold of 0.25. All visualization plots were produced using the packages Seurat, ggplot2, ComplexHeatmap, or GraphPad Prism.

### scRNA-Seq analysis – differential expression, functional enrichment

DEG analysis between groups was performed using the MAST algorithm (53) on the log-normalized RNA expression matrix using Seurat’s *FindMarkers* function with a logfc.threshold of 0. To account for unequal cell numbers between groups, cells were randomly subsampled from the larger group to match the number of cells in the smaller group prior to DEG analysis. A gene was considered significantly differentially expressed if the adjusted *P*-value was 0.05 and the |log_2_ fold change| was greater than 0.3. Enrichment analyses were performed using escape with hallmark pathways. Module scores were calculated using the Seurat *AddModuleScore* function.

### Flow cytometry for identifying γδ T cell subsets

Freshly thawed PBMC samples were rested overnight at 37°C with 5% CO_2_ in the presence of anti-CD107a BB700 (BD Biosciences, clone IP26) prior to staining with the remaining antibodies. 20 hours after thaw, anti-CD137 BUV395 (BD Biosciences, clone H4A3) was added along with brefeldin A and monensin (BD Biosciences) and kept at 37°C for another 4 hours. At the end of incubation, cells were stained with Zombie UV (BioLegend) according to manufacturer’s recommendation. Cells were then stained with unconjugated anti-TCRγδ (BD Biosciences, clone 11F2) for 20 minutes, washed, stained with secondary goat anti-mouse-Ig PE (BioLegend, clone Poly4053) for 20 minutes, and washed again. The rest of the surface antibodies were then added: anti-CD14 PE/Fire 810 (BioLegend, clone S18004B), anti-CD19 BUV805 (BD Biosciences, clone HIB19), anti-CD3 APC-R700 (BD Biosciences, clone UCHT1), anti-CD4 APC-eF780 (Invitrogen, clone RPA-T4), anti-CD8a BV650 (BioLegend, clone RPA-T8), anti-Vδ1 FITC (Invitrogen, clone TS8.2), anti-Vδ2 APC (BioLegend, clone B6), anti-Vα24-Jα18 PE/Dazzle 594 (BioLegend, clone 6B11), anti-Vα7.2 PE-Cy7 (BioLegend, clone 3C10), anti-CD161 BV421 (BioLegend, clone HP-3G10), anti-CD26 BV510 (BD Biosciences, clone L272), anti-CD16 RB545 (BD Biosciences, clone 3G8), anti-CD56 PerCP-eFluor710 (Invitrogen, clone CMSSB), anti-TCRαβ BUV563 (BD Biosciences, clone IP26), anti-CD69 BV605 (BD Biosciences, clone FN50), and anti-CD137 BUV395 again to bolster the earlier staining step. After surface staining was complete, cell membranes were rendered permeable using Cyto-Fast Fix/Perm Buffer Set (BioLegend). Intracellular staining was then performed using anti-IFNγ RB780 (BD Biosciences, clone B27), anti-IL-10 BV711 (BD Biosciences, clone JES3-9D7), anti-IL-17A BV786 (BD Biosciences, clone N49-653), anti-IL-2 BUV737 (BD Biosciences, clone MQ1-17H12), anti-TNF PerCP-Cy5.5 (Invitrogen, clone Mab11), anti-granulysin AF647 (BioLegend, clone DH2), and anti-granzyme B Pacific Blue (BioLegend, clone GB11). Samples were analyzed on a Cytek Aurora with 5L configuration. Partial data shown in Figures 5C (unstimulated).

### Quantification and Statistical Analysis

Statistical analyses were performed using Friedman test with Dunn’s multiple comparisons test, pairwise Wilcoxon rank-sum tests with Bejamini-Hochberg multiple testing correction, exact binomial test, Pearson correlation, permutation tests, two-way ANOVA with Tukey’s multiple comparisons test, or Kruskal-Wallis test with Dunn’s multiple comparisons test where appropriate (listed in the caption for each figure). Prism 10.6.0 (GraphPad) and R (v4.4.2) were used for these calculations. Values pertaining to significance are noted in the respective figure, and *P* < 0.05 was defined as statistically significant.

---

**Figure S1.**
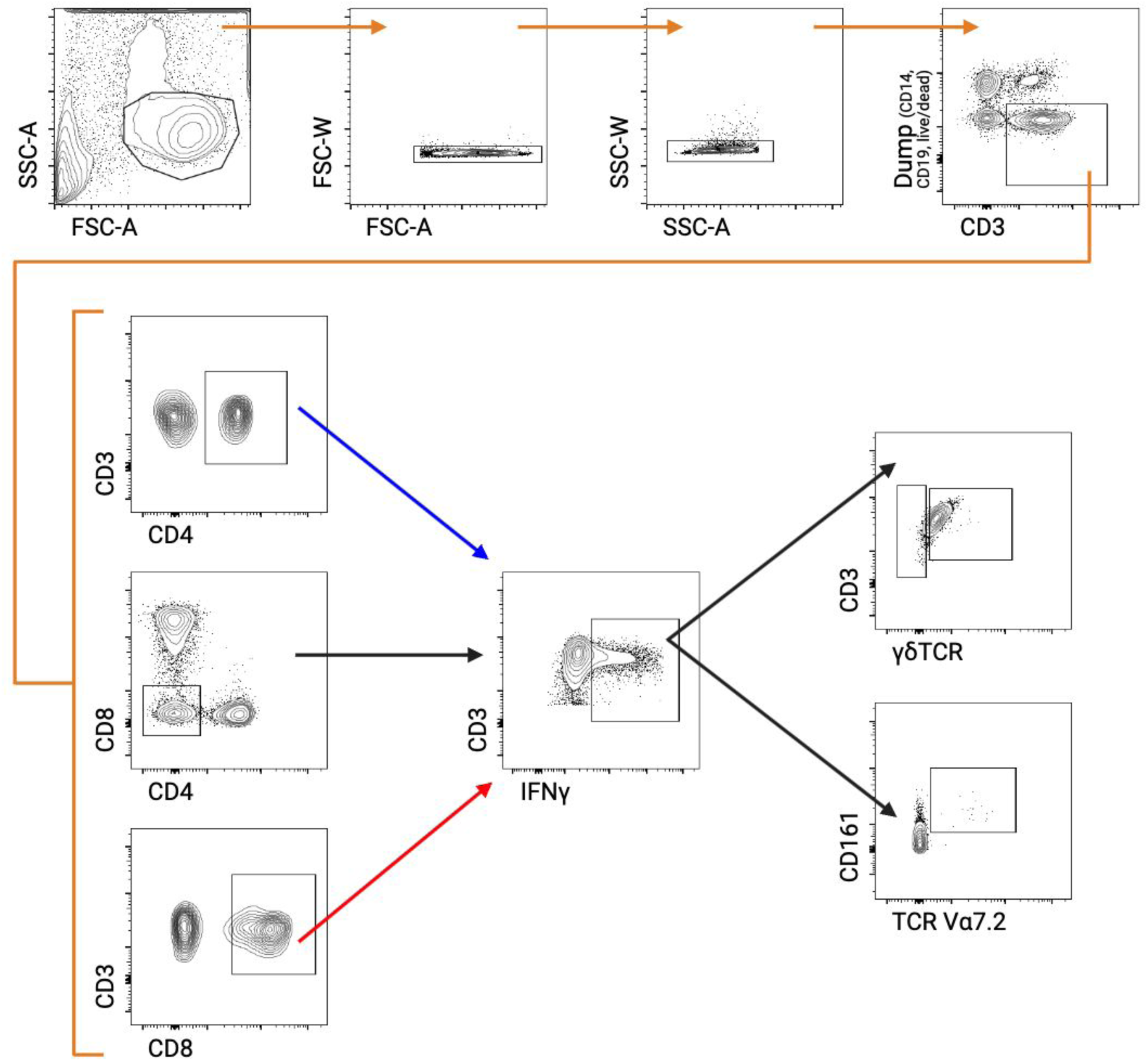
Gating strategy for analysis of IFNγ expression in response to BCG and *Mtb* peptide pool stimulation (Fig. 1).

**Figure S2.**
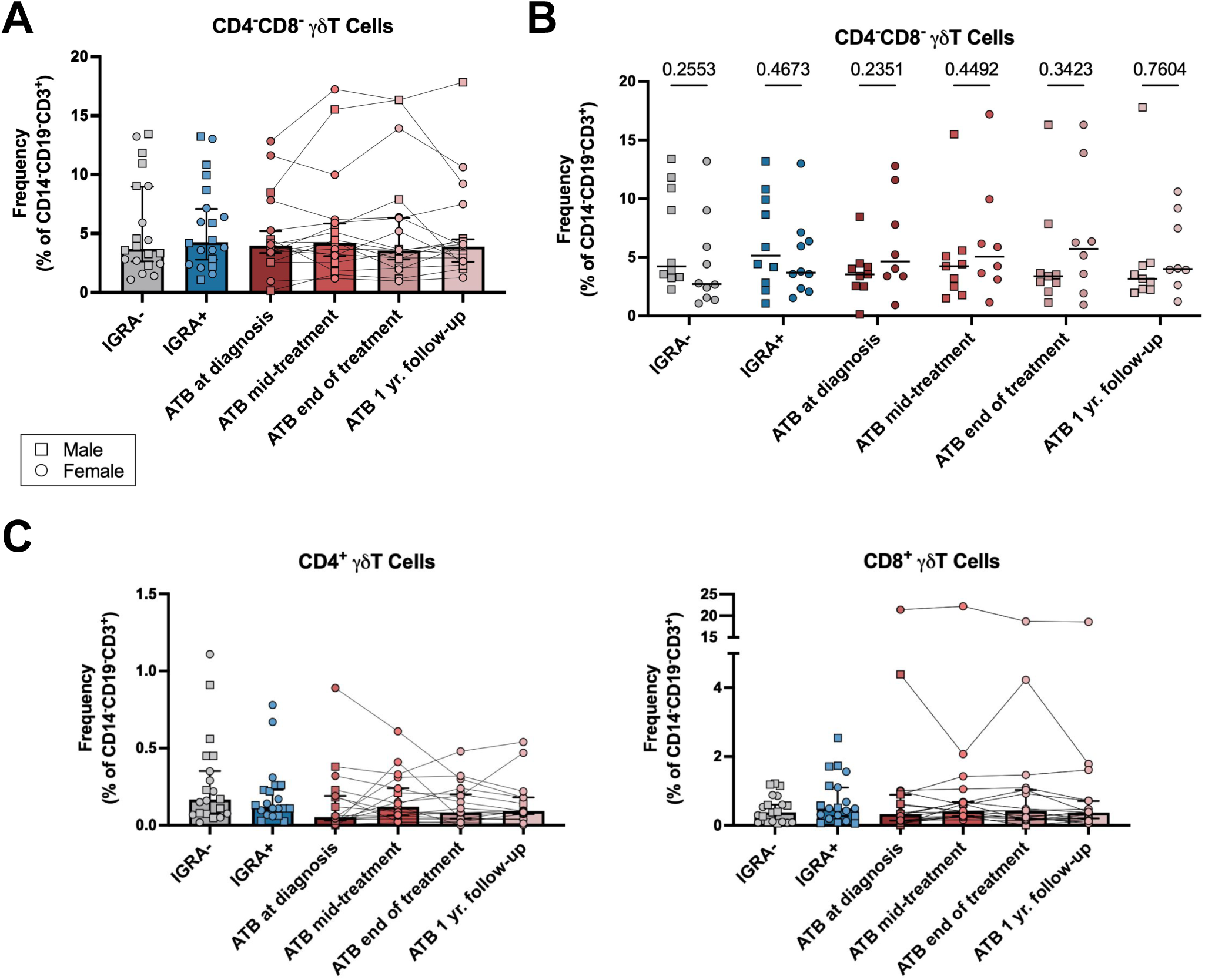
A) Frequency of CD4^-^CD8^-^ γδ T cells for each cohort measured via flow cytometry. B) Frequency of CD4^-^CD8^-^ γδ T cells for each cohort split by biological sex measured via flow cytometry. Ordinary two-way ANOVA with Tukey’s multiple comparisons test. C) Frequency of CD4^+^ (left) and CD8^+^ (right) γδ T cells for each cohort measured via flow cytometry.

**Figure S3.**
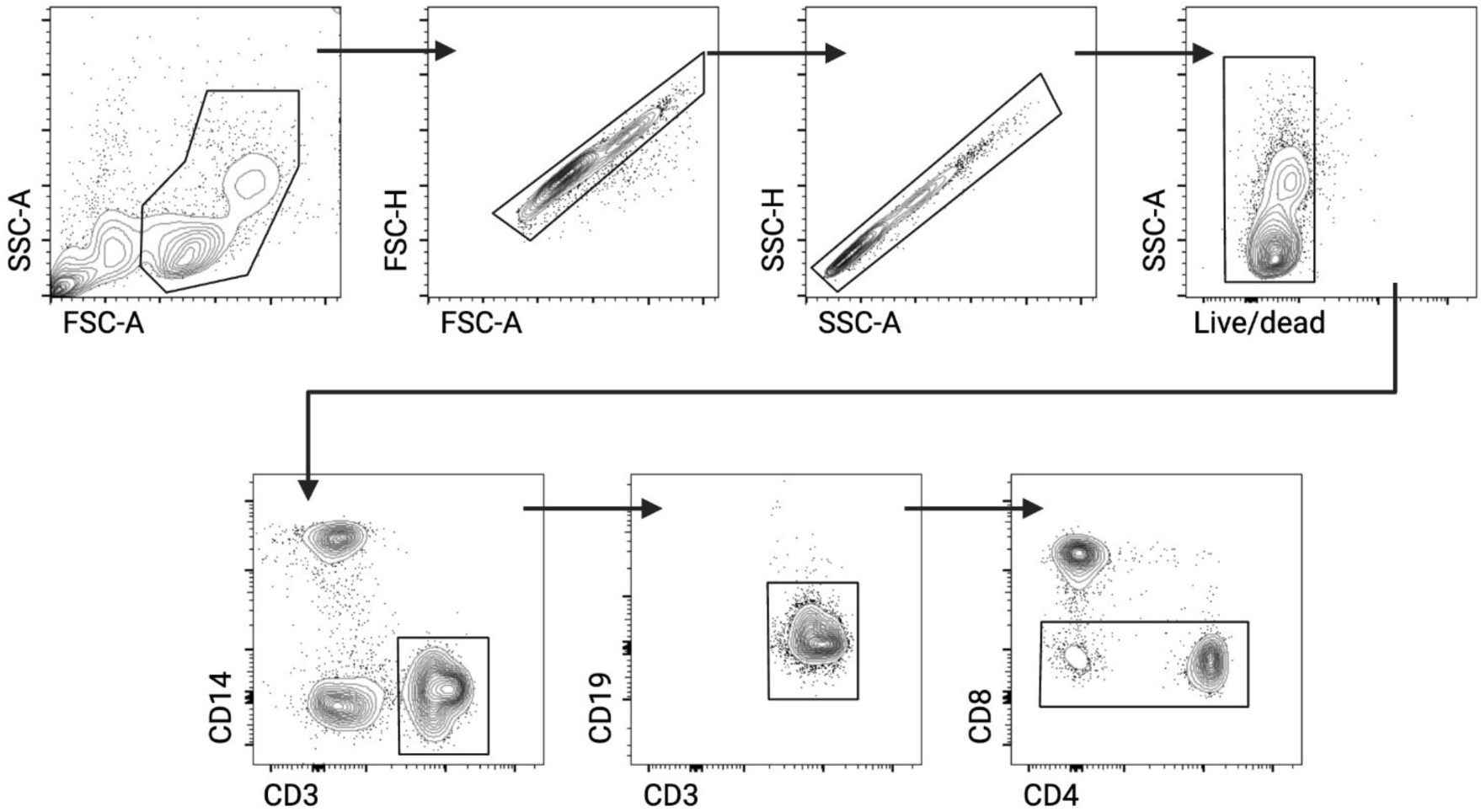
Gating strategy for fluorescence-activated sort of CD8^-^ T cells for single-cell RNA sequencing.

**Figure S4.**
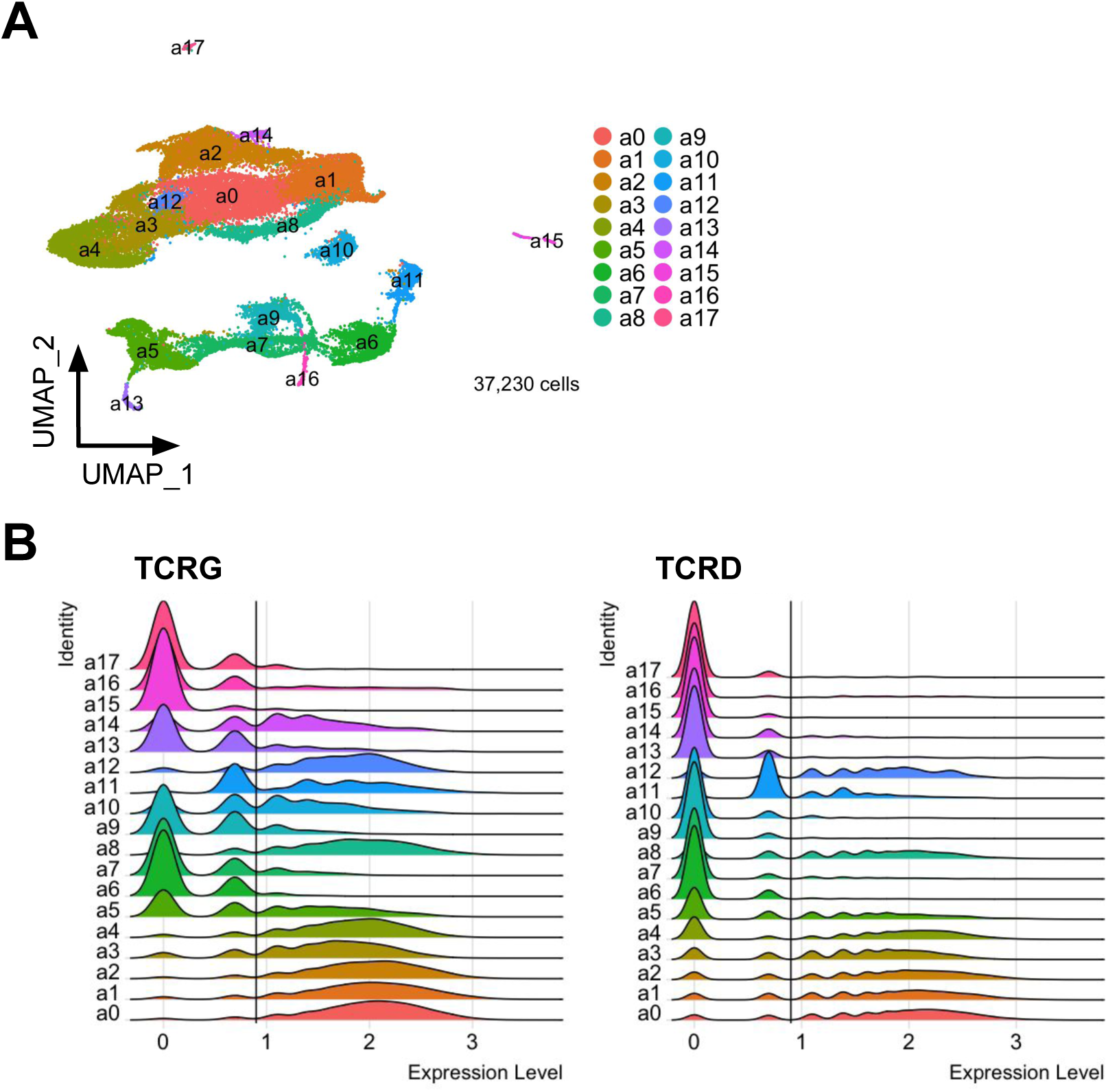
A) UMAP of all sorted CD4^-^CD8^-^ T cells. B) Expression level (SCT) of aggregated TCRG and TCRD genes per cluster as in A. Cells with expression of aggregated TCRG and TCRD > 0.9 were reclustered for further analysis (Fig. 3A).

**Figure S5.**
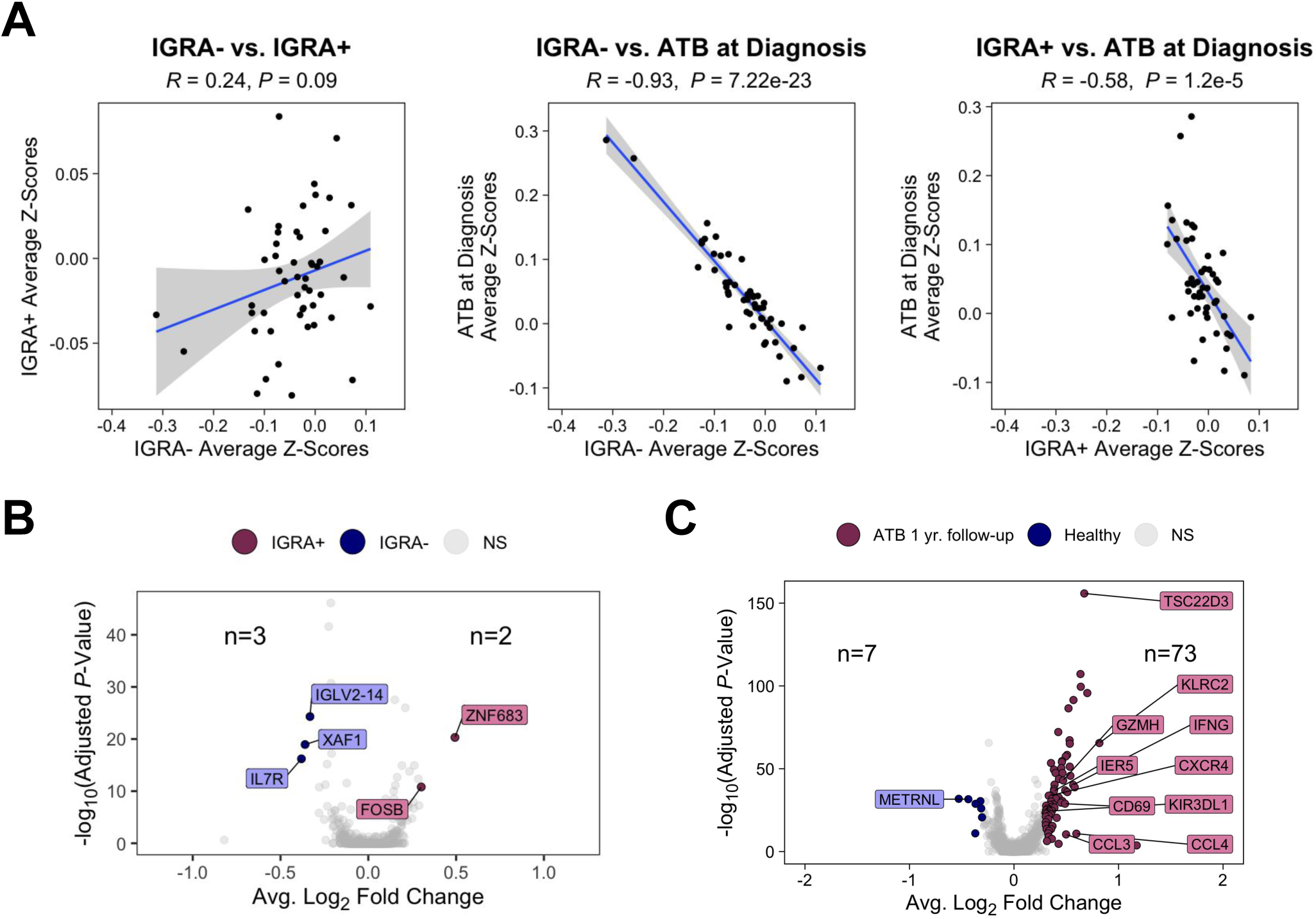
A) Pearson correlation of average GSEA Z-scores of each pathway between healthy and ATB at diagnosis cohorts. B) Differential gene expression analysis between IGRA- and IGRA+ cohorts. C) Differential gene expression analysis between ATB 1 yr. follow-up and healthy (IGRA-, IGRA+) cohorts.

**Figure S6.**
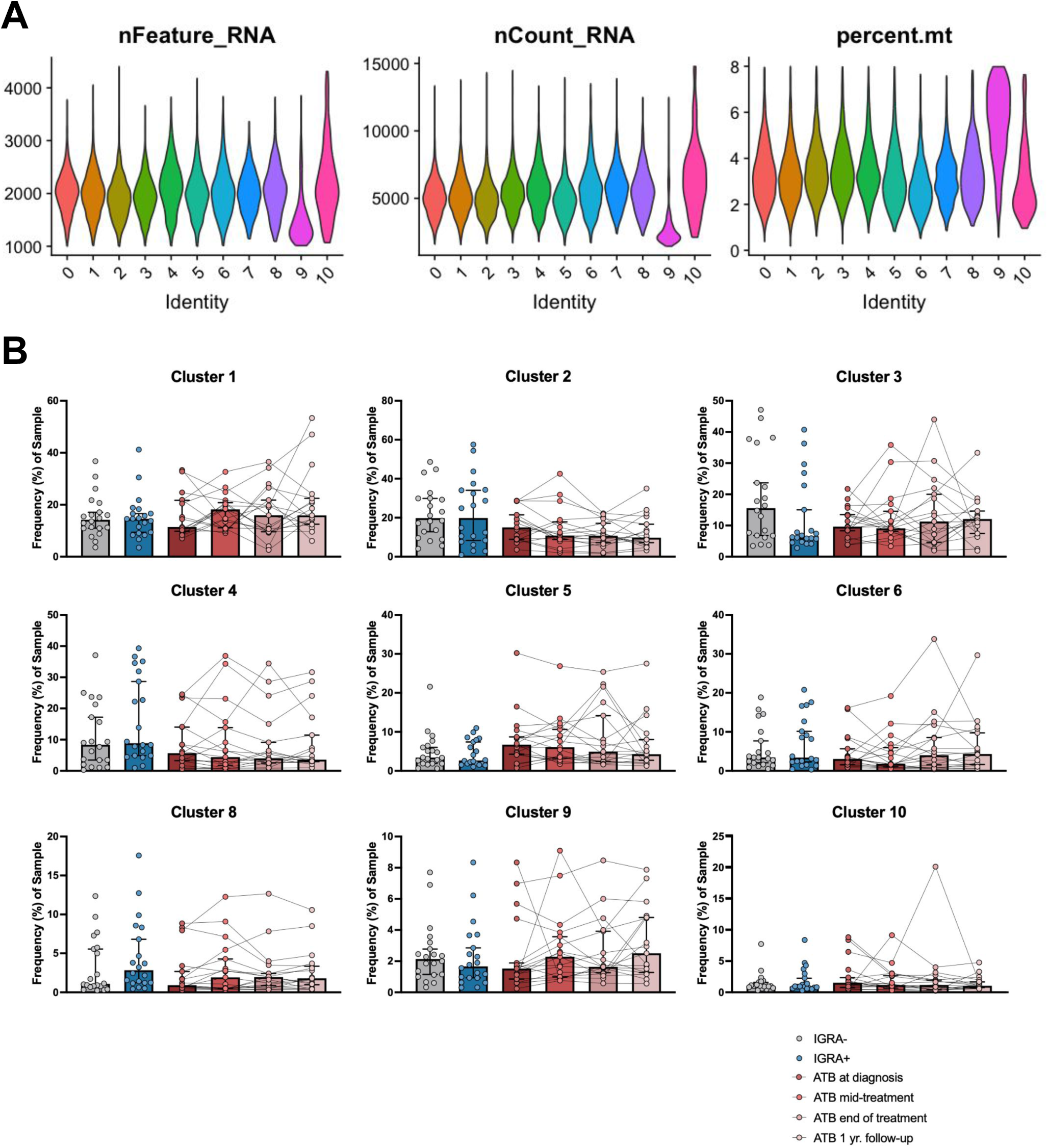
A) scRNAseq quality control results per cluster. B) Frequency of each sample found in each cluster. For unpaired analyses, Kruskal-Wallis test with Dunn’s multiple comparisons test. For paired analyses, Friedman test with Dunn’s multiple comparisons test.

